# Malaria and Burkitt’s Lymphoma: An *In Silico* Analysis of Gene Expression Links between Malaria and Burkitt’s Lymphoma and Potential Anticancer Activity of Artemisinin Derivatives

**DOI:** 10.1101/356782

**Authors:** Inas Elsayed, Mutaz Amin, Muzamil Mahdi Abdel Hamid, Xiaosheng Wang, Mie Rizig

**Author notes:** Corresponding author: Inas Elsayed, Faculty of Pharmacy, University of Gezira. P. O. Box 20. I.E: Inas Elsayed. M. R: Mie Rizig.

## Abstract

**Background:** Burkitt’s lymphoma (BL) is an aggressive form of B-cell non-Hodgkin lymphoma. Endemic subtype of the disease showed a remarkable statistical and epidemiological association with malaria infection. Despite the numerous studies performed to explain this association; molecular mechanisms underlie such coincidence still remain unclear. Dissecting molecular mechanisms which link Malaria infection and Burkitt’s lymphoma can provide insights about reported anticancer action of certain antimalarial drugs, namely artemisinin derivatives.

**Methods:** Here we applied an integrative approach to investigate for potential links between malaria infection and endemic Burkitt’s lymphoma regarding their gene expression, and further explore common molecular mechanisms through which artemisinin compounds might act in endemic Burkitt’s lymphoma. Using gene expression data of malaria (Plasmodium falciparum infected erythroblasts) and endemic Burkitt’s lymphoma from Gene Expression Omnibus database, expression patterns in the two conditions were examined through clustering analysis using Self Organizing Maps, and then by significance testing of differentially expressed genes in each condition followed by Functional annotation using Gene Ontology clustering and Pathways analysis.

**Results:** Clustering analysis identified a significant overlap between the expression patterns in endemic Burkitt’s lymphoma and Plasmodium falciparum infected cells. Four out of the 12 identified clusters contained genes with similar expression patterns in both conditions. Differential expression analysis identified 1689 genes as significantly differentially expressed in endemic Burkitt’s lymphoma and 405 in malaria. Those genes were found to be related to important Gene Ontology terms and pathways. Interestingly 65% of the identified pathways in Malaria were overlapped with those identified in endemic Burkitt’s lymphoma. Several of these pathways reported to be related to actions of artemisinin derivatives.

**Conclusion:** Our In-silico analysis showed a significant molecular convergence between endemic Burkitt’s lymphoma and malaria. A number of 43pathways which demonstrated enrichment in tumour were shared with *Plasmodium falciparum* infected erythrocytes. Such pathways represent potential targets for antimalarial drugs to exert therapeutic effects in such malignancy.

## Background

Burkitt’s lymphoma (BL) is a highly aggressive, monoclonal, B-cell derived, non-Hodgkin lymphoma [1, 2]. According to cells’ morphology, phenotype and genetic profile BL is generally classified as a germinal center neoplasm [1]. Currently, three clinical /epidemiological variants of the disease have been defined according to WHO classification −2008; Endemic “African” (eBL), sporadic (sBL), and acquired immunodeficiency syndrome (AIDS)–related BL. The endemic type of Burkitt’s lymphoma is invariably linked to EBV Pre-infection which proved to play major role in disease pathogenesis [2, 3, 4]. In addition, reports revealed a curious association between eBL incidence and rates of ***Plasmodium falciparum*** *(P. falciparum)* malaria infection. Despite the well-established role of EBV infection, and the suspected implication of malaria, in the genesis of eBL, very little research has examined the interaction between the two pathogens. Both agents actually cause B cell hyperplasia, which is considered almost certainly an essential component of lymphomagenesis in BL [2, 5, 6].

Studies carried out to clarify the role of *P. falciparum* malaria in the eBL etiology came out with two proposed theories. The first theory implies that *P. falciparum* malaria induces polyclonal B-cell expansion and reactivation of lytic EBV infection leading to the expansion of latently infected B-cells and hence, raises the likelihood of c-myc translocation. The second theory states that EBV-specific T-cell immunity is impaired during *P. falciparum* malaria co-infection due – or may be in consequence-to EBV replication, which eventually leads to loss of viral control [7]. However, The impact of malaria on the control of EBV persistence seems to be evident only before immunity to malaria is fully acquired, since adults in such endemic areas do not have detectable EBV DNA in the plasma in contrast to children living in the same area [1] Besides, eBL is a childhood tumour with a peak age incidence between 5 and 8 years; an age at which malaria infection may impair the virus-host balance [6].

On the other side applying malaria control programs using antimalarial drugs was followed by a considerable reduction in incidence of eBL. Artemisinin and its derivatives such as artesunate (ART) were main players in such programs. Artemisinin and its related compounds have been distinguished as a new generation of antimalarial drugs with the advantage of activity against multidrug-resistant *Plasmodium falciparum* and *Plasmodium vivax* strains, that’s not compromised by obvious adverse reactions. Recently, the antimalarial artemisinin drugs – especially ART-also been reported to have anti-tumour activity in several cancer cell lines including chemotherapy resistant ones [8, 9, 10].

Hence, elucidation of common pathways between malaria infection and eBL tumour can help explaining the suggested anti-tumorous activity of artemisinin. This study aimed to elucidate the gene expression links between eBL and malaria infected erythroblasts, and the pathways shared between them in order to find a common ground between the two diseases and possibly explain the role of artemisinin derivatives as a potential anticancer agent in eBL.

## Materials and methods

### Data recruitment

Gene expression data was recruited through the gene expression omnibus (GEO)[11, 12]. GEO data series was searched for all the studies under the terms (Malaria) and (Burkitt’s lymphoma). The recruited studies were further compared and suitable studies were chosen on the basis of their matched experimental conditions and the type of arrays used. For the sake of safe data comparability among different samples we considered studies using the same microarray platform. All samples used in the analysis were hybridized to the Affymetrix Human Genome U133 plus 2.0 Array [HG-U133_Plus_2] which contains over 47,000 transcripts and were prepared following the standard Affymetrix protocol. Description of the HG-U133_Plus_2 array can be viewed on the Affymetrix website [13]. Affymetrix^®^ platform has been preferred for the reasons of considerable precision and range of gene expression magnitude as well as excellent annotation that keeps concordance of platform output when used in different studies.

### Statistical analysis

We applied two types of analysis: (1) Clustering technique using Self-Organising Maps (SOM) to examine the expression patterns (i.e. the degree of the differences and the similarities) between the eBL and malaria samples, and (2) Significant analysis using T test to find the significant genes within each group (i.e. eBL and malaria) compared to control.

#### (A) Clustering analysis using Self Organising Maps (SOM)

Three biological replicates per each condition (3 samples of malaria infected erythrocytes and 3 samples of eBL tumours) were recruited for this analysis as described above. Details of these samples can be attained through GEO viewer using the following accession numbers: {GEO: GSM656452, GSM656453, GSM656454 for BL replicates and GSM611252, GSM611253, GSM611254} for Malaria replicates [14, 15]. The intensity Cell files were downloaded to Gene-spring software for further statistical analysis [16]. Intensity signals were normalized using the *Robust Multi-array* Analysis Summarization method (RMA) [17]. The correlations of the normalized data were tested using Whisker Box plots. Normalized values were then subjected to clustering analysis using self-organizing maps (SOM). SOM was built using the default setting within Gene Spring software which applies the Euclidean algorithm with the following criteria: maximum number of iteration is 50; neighborhood radius of 5 and a Grid size was 3×4.

#### (B) Statistical analysis using significance testing

Two studies (one for malaria and one for BL) were chosen again as described previously “refer to table 1 for studies’ description”. Five biological replicates of eBL samples were compared to five replicates of control “normal Centeroblasts”. Details of samples can be accessed through GEO viewer using the following accession numbers; {GEO: GSM312851, GSM312853, GSM312854, GSM312856, GSM312857} for eBL samples and {GEO: GSM312937, GSM312938, GSM312939, GSM312940, GSM312941} for BL controls. Statistical analysis was performed using GEO2R which applies; Linear Models for Microarray Analysis (LIMMA) R packages from the Bioconductor project [18, 19]. Data were normalized using variance stabilization method, and value distributions of normalized intensities were compared using box plots (Fig.1,2, and 3) [20]. Up/down regulated genes with a significant t test;(P value < 0.05) after applying Bonferroni correction for multiple testing; have been identified. The same analysis has been applied to malaria samples where three replicates of malaria “infected erythroblasts” and three control samples “uninfected erythroblasts” used. Detailed information about samples can be attained through GEO accession viewer using the accession numbers: {GEO: GSM611252, GSM611253, GSM611254} for Malaria samples, and {GEO: <u>GSM611247</u>, <u>GSM611248</u>, <u>GSM611249</u>} for malaria control. Full description of the two studies can be accessed through the GEO database using provided accession numbers [21, 15].

**Table 1:**
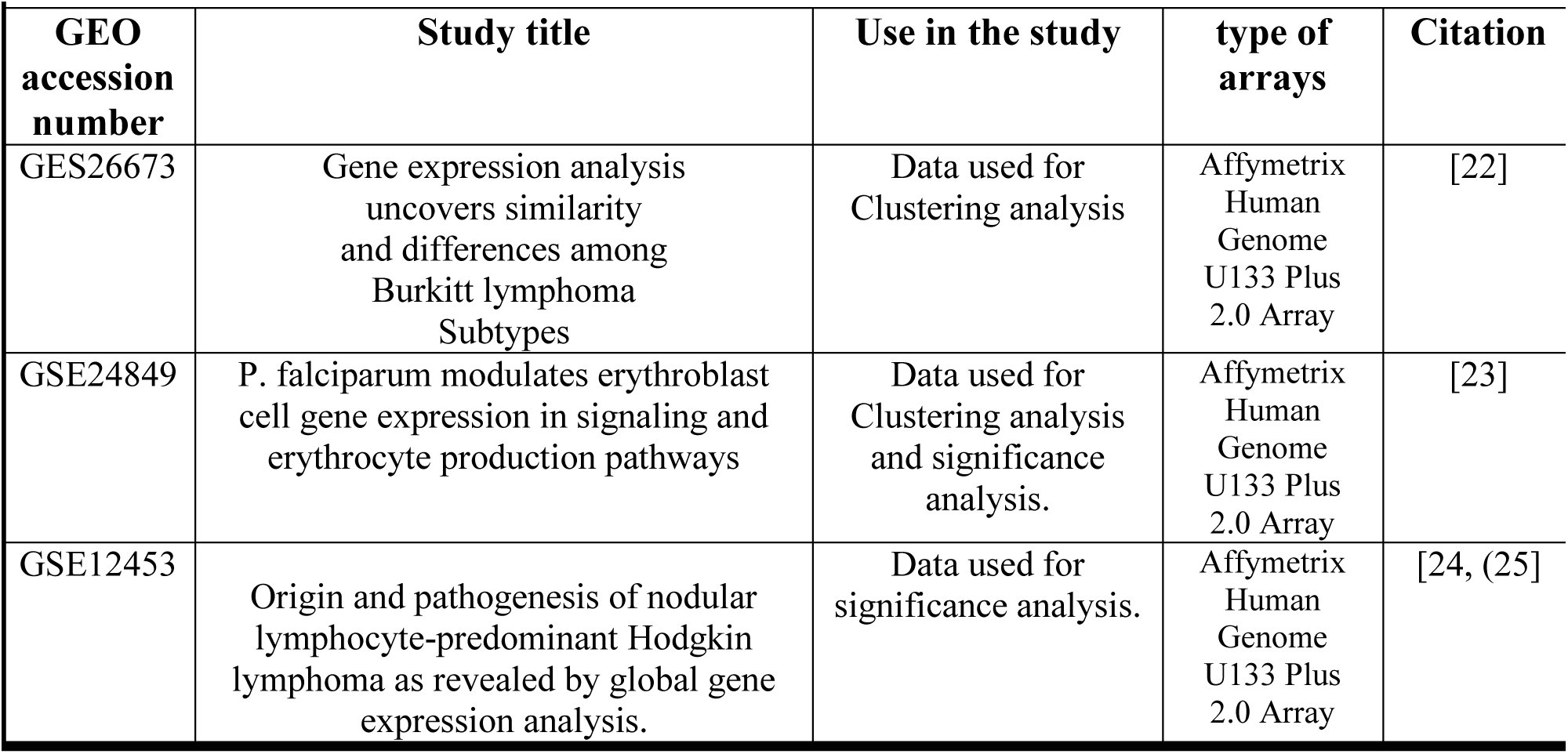
Description of the studies:

| GEO accession number | Study title | Use in the study | type of arrays | Citation |
| --- | --- | --- | --- | --- |
| GES26673 | Gene expression analysis uncovers similarity and differences among Burkitt lymphoma Subtypes | Data used for Clustering analysis | Affymetrix Human Genome U133 Plus 2.0 Array | [22] |
| GSE24849 | P. falciparum modulates erythroblast cell gene expression in signaling and erythrocyte production pathways | Data used for Clustering analysis and significance analysis. | Affymetrix Human Genome U133 Plus 2.0 Array | [23] |
| GSE12453 | Origin and pathogenesis of nodular lymphocyte-predominant Hodgkin lymphoma as revealed by global gene expression analysis. | Data used for significance analysis. | Affymetrix Human Genome U133 Plus 2.0 Array | [24, (25)] |

**Figure (1.a).**
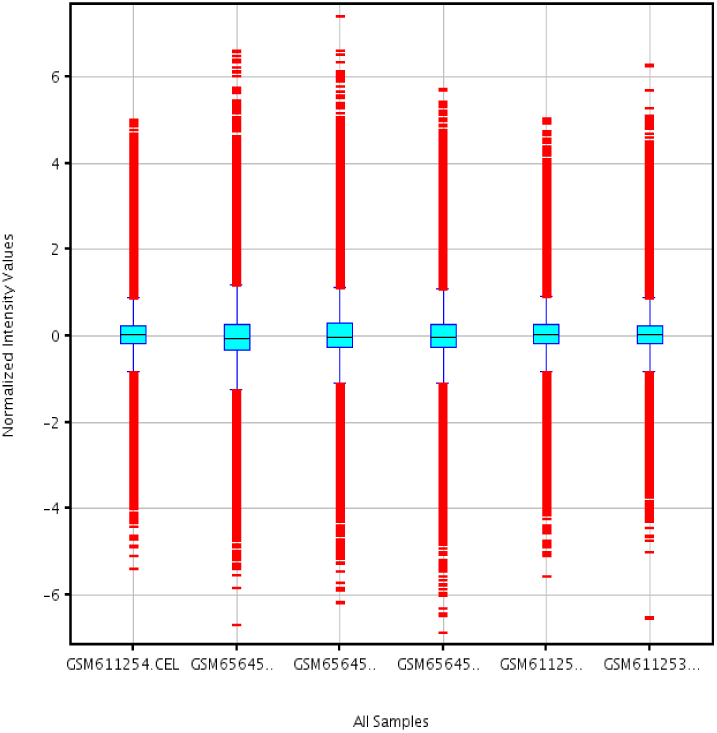
Box plot of the normalized intensities-eBL,GES26673. Figure represents a distribution of gene expression intensities of eBL samples; {GEO: GES26673} after normalization.

**Figure (1.b).**
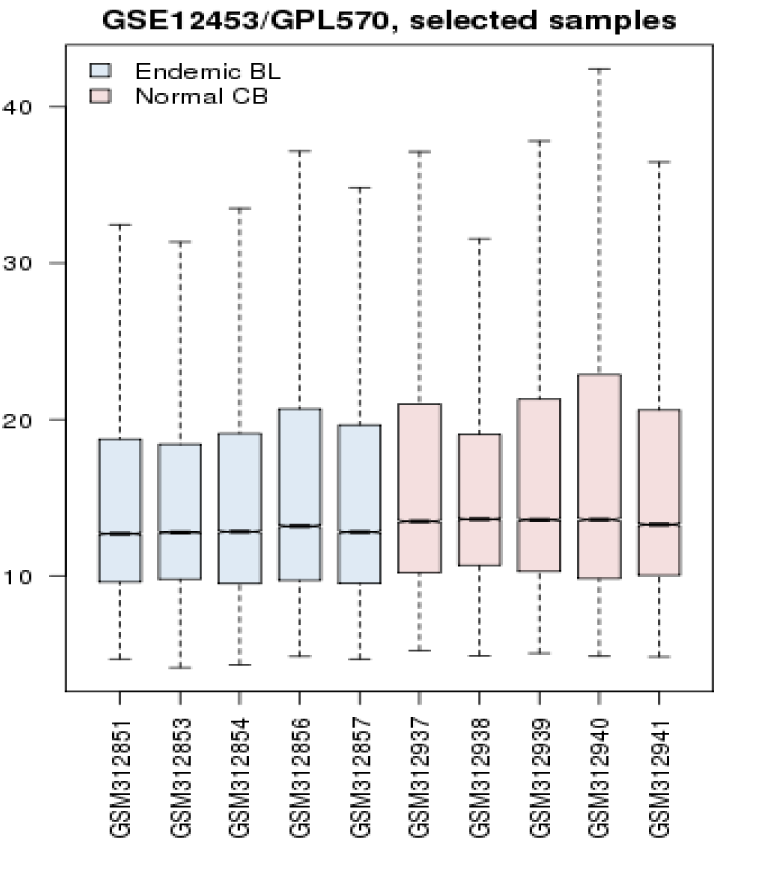
Box plot of the normalized intensities- BL, GSE12453. Figure represents a distribution of gene expression intensities of eBL samples; {GEO: GSE12453} after normalization.

**Figure (1.c).**
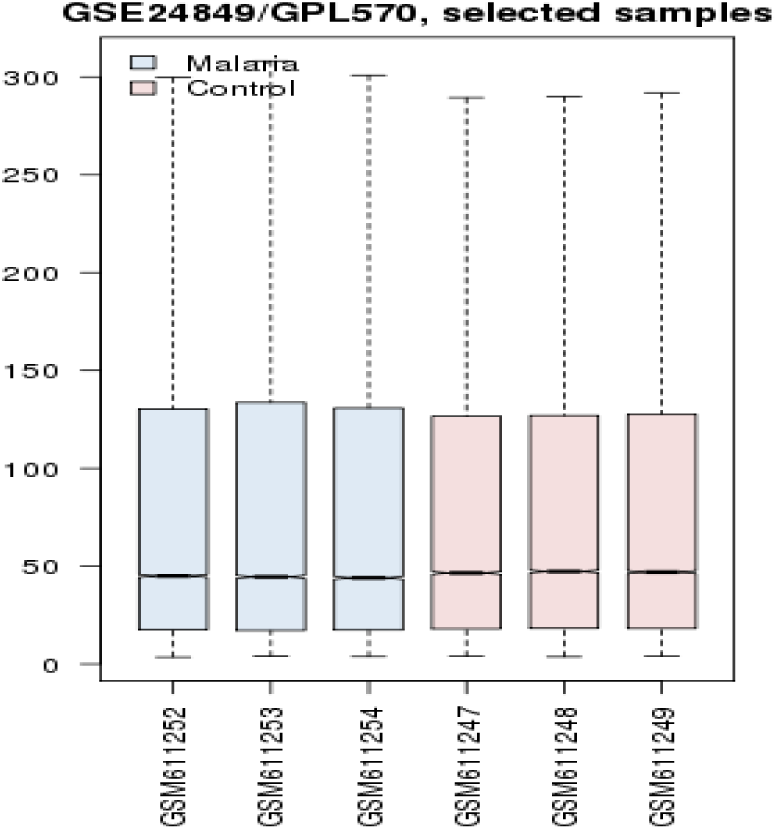
Box plot of the normalized intensities- Malaria, GSE24849. Figure represents a distribution of gene expression intensities of malaria samples; {GEO: GSE24849} after normalization.

### Data mining (Functional annotation using Gene Ontology and Pathways analysis)

Genes verified as significantly differentially expressed, t test (p value < 0.05), in each condition were subjected to functional annotation clustering and pathways analysis using functional classification tool of Database for Annotation, Visualization and Integrated Discovery database (DAVID) (http://david.abcc.ncifcrf.gov/). Significant Gene ontology (GO)^a^ clusters “using Bonferroni multi-test correction” and pathways “using KEGG pathway option ^b^” have been identified in eBL as well as malaria state.

## Results

### GEO search

Interrogating GEO data series for the terms malaria and BL resulted in 20 studies for malaria and 19 studies for lymphoma. Three of them have been used in this study; two for eBL and one for malaria. Full description of the studies which are included in the analysis within this study is included in table 1.

### Data normalization

Box plots of normalized intensities of all the samples are shown in fig. (1, 2, and 3).

### Clustering analysis using Self-Organizing Maps (SOM)

SOM were used to analyze the differences and similarities within the gene expression patterns between malaria and eBL. Analysis produced 11 clusters of genes in both malaria and eBL. Eventually, defined patterns of behavior were observed among clusters within the two conditions. These include clusters of genes which are down regulated in eBL and up regulated in malaria (namely cluster 0, 1, and8), genes which are Up-regulated in eBL and down regulated in malaria (namely cluster 3, 7, 10, and 11), and finally, Genes with similar expression signals in eBL and malaria (namely cluster 4, 7, 6, and 9). Figure 4 illustrates the patterns of expression of these clusters in both conditions. The total number of genes with a similar behavior was 30,532 out of 47,000 (i.e.; about 65%). List of those similarly expressed genes is provided in supplementary tables S1, S2, S3, and S4. Table 2 summarizes the SOM findings. These findings hint towards a similarity within the expression patterns and perhaps the molecular mechanisms between the two tested conditions. To investigate such possibility we proceeded to test the differential expression patterns and pathways analysis in each condition.

**Figure 4:**
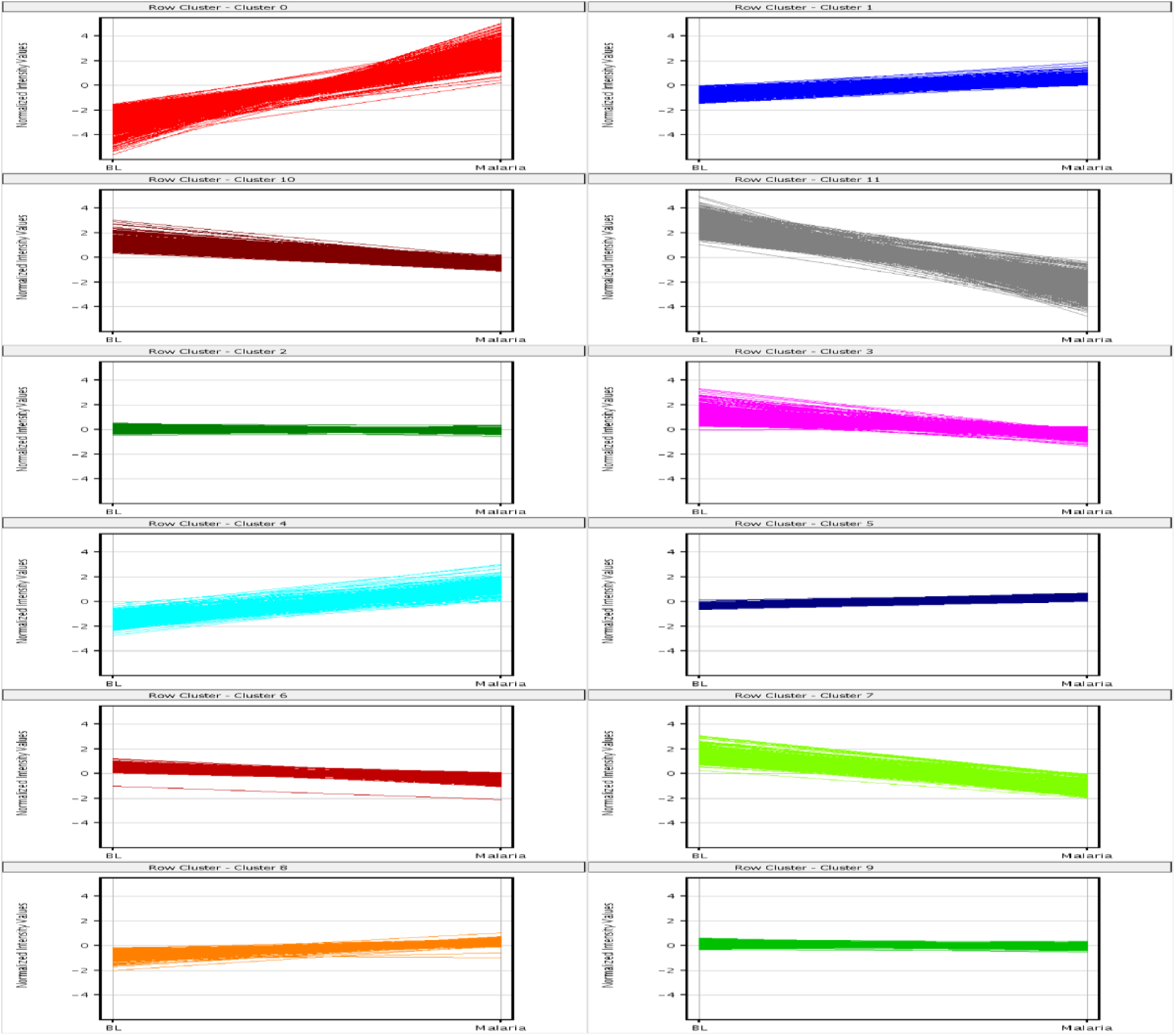
(4) 3×4 Self organising maps for Malaria and BL samples. Figure illustrates the patterns of gene expression in both eBL and malaria conditions. These patterns were clustered mainly around five groups. Clusters of gene which are down regulated in BL and up regulated in Malaria (namely cluster 0 and 1). Genes which are Up-regulated in BL and down regulated malaria (namely cluster 3, 7, 10, and 11). Genes which are down regulated in BL and up-regulated in Malaria (cluster 8). Finally Genes with similar expression signals in eBL and malaria (namely cluster 4, 7, 6, and 9).

**Table 2:**
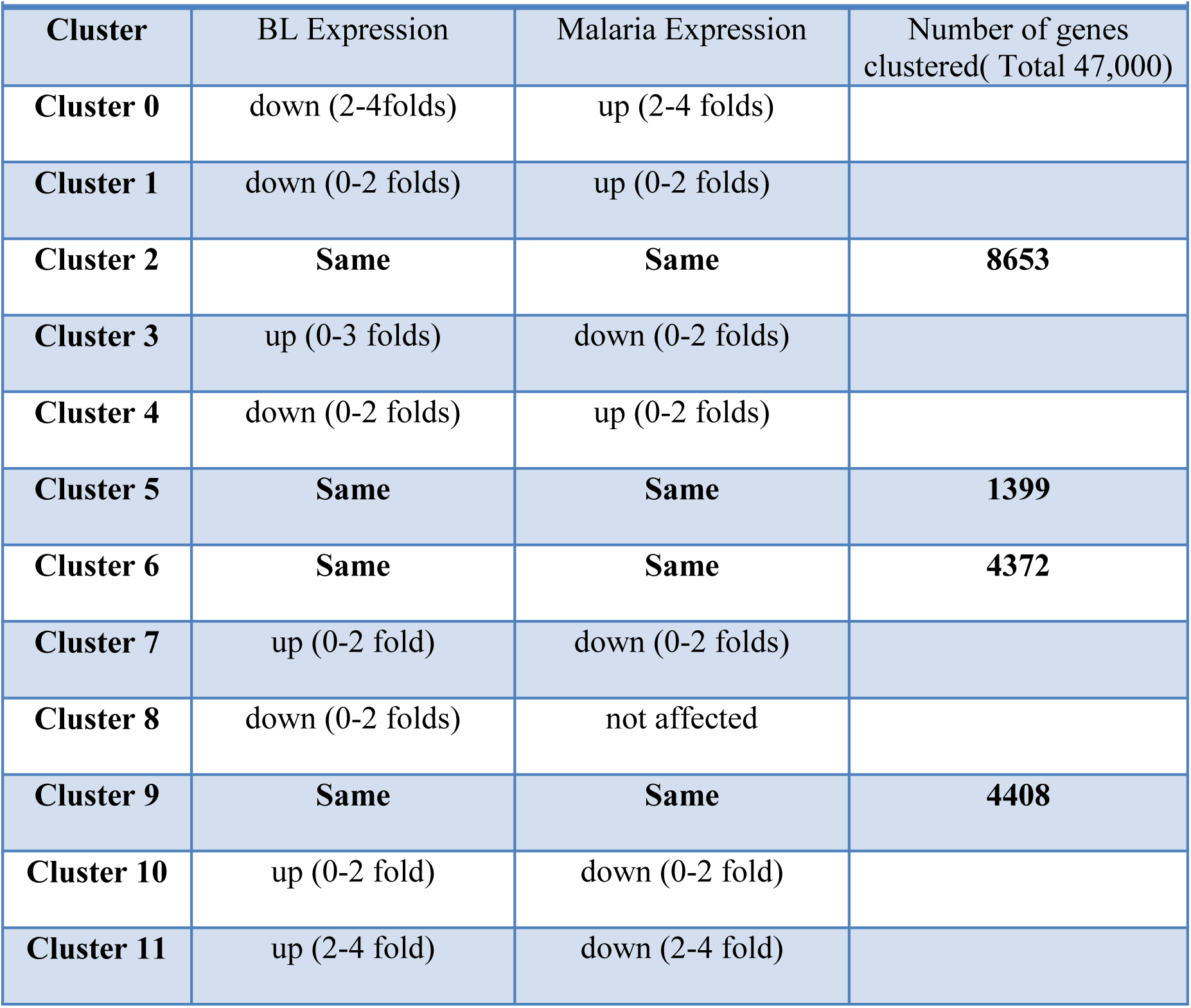
SOM clusters of gene expression in eBL cells and malaria in vitro infected erythroblasts from GEO database

|  |  |  |  |
| --- | --- | --- | --- |
| Cluster | BL Expression | Malaria Expression | Number of genes clustered( Total 47,000) |
| **Cluster 0** | down (2-4folds) | up (2-4 folds) |  |
| **Cluster 1** | down (0-2 folds) | up (0-2 folds) |  |
| **Cluster 2** | **Same** | **Same** | **8653** |
| **Cluster 3** | up (0-3 folds) | down (0-2 folds) |  |
| **Cluster 4** | down (0-2 folds) | up (0-2 folds) |  |
| **Cluster 5** | **Same** | **Same** | **1399** |
| **Cluster 6** | **Same** | **Same** | **4372** |
| **Cluster 7** | up (0-2 fold) | down (0-2 folds) |  |
| **Cluster 8** | down (0-2 folds) | not affected |  |
| **Cluster 9** | **Same** | **Same** | **4408** |
| **Cluster 10** | up (0-2 fold) | down (0-2 fold) |  |
| **Cluster 11** | up (2-4 fold) | down (2-4 fold) |  |

### Differential expression analysis

Differential expression analysis of eBL and malaria datasets using GEO2R tool produced 1689 differentially expressed genes for eBL and 405 for malaria (supplementary tables S5, and S6). Each set of differentially expressed genes has been analyzed using DAVID database functional annotations tools for identification of pathways and functional clusters the genes are most likely to be implicated in. Analysis resulted in 320 and 74 functional clusters (supplementary tables S9, and S10), in addition to 353 and 66 pathways analysis records (supplementary tables S7, and S8) for eBL and malaria successively. Interestingly, about 65% of malaria pathways found to be shared with eBL; table 3 lists pathways found to be common between malaria and eBL.

**Table 3:**
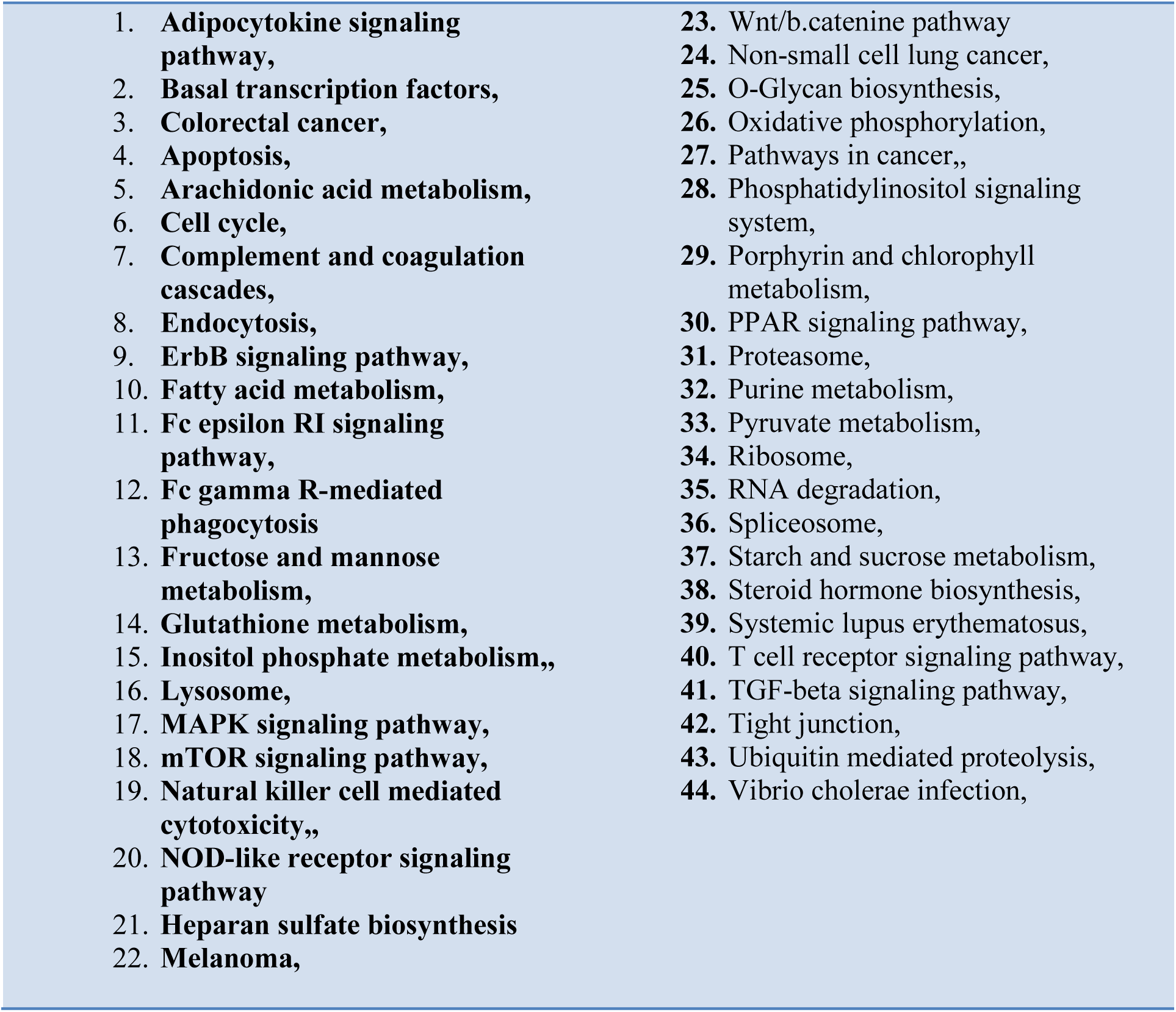
Common pathways between eBL cells and malaria in vitro infected erythroblasts from GEO database

|  |  |
| --- | --- |
| 1. **Adipocytokine signaling pathway,** | 23. Wnt/b.catenine pathway |
| 2. **Basal transcription factors,** | 24. Non-small cell lung cancer, |
| 3. **Colorectal cancer,** | 25. O-Glycan biosynthesis, |
| 4. **Apoptosis,** | 26. Oxidative phosphorylation, |
| 5. **Arachidonic acid metabolism,** | 27. Pathways in cancer,, |
| 6. **Cell cycle,** | 28. Phosphatidylinositol signaling system, |
| 7. **Complement and coagulation cascades,** | 29. Porphyrin and chlorophyll metabolism, |
| 8. **Endocytosis,** | 30. PPAR signaling pathway, |
| 9. **ErbB signaling pathway,** | 31. Proteasome, |
| 10. **Fatty acid metabolism,** | 32. Purine metabolism, |
| 11. **Fc epsilon RI signaling pathway,** | 33. Pyruvate metabolism, |
| 12. **Fc gamma R-mediated phagocytosis** | 34. Ribosome, |
| 13. **Fructose and mannose metabolism,** | 35. RNA degradation, |
| 14. **Glutathione metabolism,** | 36. Spliceosome, |
| 15. **Inositol phosphate metabolism,,** | 37. Starch and sucrose metabolism, |
| 16. **Lysosome,** | 38. Steroid hormone biosynthesis, |
| 17. **MAPK signaling pathway,** | 39. Systemic lupus erythematosus, |
| 18. **mTOR signaling pathway,** | 40. T cell receptor signaling pathway, |
| 19. **Natural killer cell mediated cytotoxicity,,** | 41. TGF-beta signaling pathway, |
| 20. **NOD-like receptor signaling pathway** | 42. Tight junction, |
| 21. **Heparan sulfate biosynthesis** | 43. Ubiquitin mediated proteolysis, |
| 22. **Melanoma,** | 44. Vibrio cholerae infection, |

## Discussion

Malaria infection has been recurrently linked to endemic form of BL. Reported association of the two diseases suggested a possible implication of malaria infection in the pathogenesis of eBL [6]. Through analysis of their expression profiles with unsupervised clustering analysis, the SOM algorithm showed a significant overlap in the gene expression patterns between the two diseases. This was demonstrated by the considerable number of the genes (30,532 genes) clustered within groups with similarly expressed signals between the two conditions. This finding was quite encouraging as it reinforced the advocate for malaria and BL molecular convergence. Such significant overlap regarding gene expression patterns of eBL and malaria suggests that antimalarial drugs which possess anticancer potentials might act through mechanisms and pathways that are common between the two diseases.

To indicate common pathological mechanisms between the two diseases we proceeded to perform a differential expression analysis in order to point out genes that are more likely to be involved in disease pathogenesis. A number of transcripts were identified as significantly differentially expressed in each condition compared to control. The functional annotations of these genes revealed a significant number of interesting Go terms and pathways related to each disease.

The most enriched Go clusters of eBL clearly reflected the escalated activity of c-myc onco-protein. This was demonstrated by the enhanced protein synthesis “cluster1”, stimulation of numerous transcription factors “clusters 2,9,10” and the subsequent protein-DNA and protein-protein interaction “clusters 2,6”, which lead eventually to B-lymphocytes activation, growth promotion, and proliferation “cluster 11” [26].

While the Go clusters in malaria on the other side expressed the growth and multiplication activity of the parasite through expression of structural proteins “clusters 1,2,13”, transcription regulators and DNA-binding proteins “clusters 3,4,6,11”, and DNA replication machinery “clusters 21,61”. Interestingly, a conflict in gene expression has been detected here as both genes which are inhibitory and those which induce apoptosis found to be differentially expressed suggesting a mechanism of the parasite activated to overcome suicidal attempts of the infected blood cell.

Interestingly, one of the enriched GO clusters of malaria, GO cluster 63, is involved in lymphocytes development and activation. We suspect that this might have an implication in the pathogenesis of eBL through the reactivation of latently EBV-infected B lymphocytes. Reactivation of latent virus to the lytic form with the subsequent expression of viral oncoproteins is claimed to be the principal role that malaria infection plays in promoting development of eBL [7].

In addition, pathways analysis revealed 43 shared KEGG pathways between the two conditions (Table.3). **Oxidative phosphorylation**, **mitogen-activated protein kinas (MAPK) signaling**, **Cell cycle**, **Endocytosis**, **RNA degradation**, **Spliceosome**, **Ubiquitin mediated proteolysis**, **Wnt/b.catenine pathway**, and **mTOR signaling pathways** are the most represented pathways within the group of differentially expressed genes in both diseases. It worths to note that a number of these common pathways have been examined in previous studies for potential pharmacological targets of artemisinin and its derivatives using variable cancer cell lines.

Artemisinin molecule contains an endoperoxide bridge that reacts with an iron atom -or reduced heme-to form free radicals “alkylating agents”, which causes macromolecular damage and cell death. Such proposed mechanism reasonably explains the high selective toxicity index of artemisinin derivatives which is due to higher ion uptake by actively proliferating tumor cells compared to normal cells. However the exact molecular mechanism and targets of artemisinin cytotoxicity remain to be illustrated [8, 10].

As anti-cancers, artemisinin and its derivatives reported to exert anti proliferative effects at low doses while inducing apoptic “or in some cases necrotic” response in higher doses. Antiproliferative activity of the drugs detected to be mediated through repressing the expression of cell cycle machinery namely Cyclines and Cycline dependent Kinases (CDKs). Apoptosis on the other hand found to be induced through modulating the ratio of the proapoptic to the antiapoptic proteins for the favor of the later ones. According to our analysis, genes of the mentioned pathways were differentially expressed in eBL cells which implies that artemisinin compounds might exert therapeutic activity on tumour through such pathways [27].

In addition, one of the most interesting findings is the activity of the Wnt/b-catenin signaling pathway in both conditions. the Wnt/b-catenin is a powerful signaling pathway that plays a crucial role in cell fate determination, survival, proliferation and movement in variety of tissues; in both malaria and BL [28]. Artemisnin derivative; artesunate, has been reported to act through regulation of Wnt/b-catenin related genes expression in human colorectal carcinoma [28]. Thus, reported activity of this pathway in eBL indicates that artesunate might be equivalently effective in controlling eBL through inhibition of such pathway.

Moreover, artesunate can also act against BL tumour through alternative mechanism by stimulating Transforming Growth Factor-βeta (TGF-β) pathway; a response proved to induce apoptosis in Myc-transformed neoplastic cells [29].

Also, artemisinin derivative “Dihydroartemisinin” proved to inhibit the Mammalian Target of Rapamycin (mTOR) signaling pathways which has a remarkable role in tumour cells survival, proliferation, metastasis and angiogenesis [30]. Genes within mTOR pathway were differentially expressed in both eBL and Malaria. This might support the claim that artemisinin control the aggressive tumor activity through (mTOR) signaling [30].

It is important to mention that though many of the Malaria/eBL common pathways identified in the study were reported to have well established implication in pathogenesis of eBL, some might require to be studied within the contest of tumor micro environment. Example of those is the Complement and coagulation cascades Pathway; which noted to have a dual action in tumor pathogenesis. The pathway may act as negative regulator preventing tumor development through immune-surveillance. While in the context of chronic inflammation pathway found to promote tumor growth. Balance between the two actions is majorly dictated by immunomodulators produced in tumor microenvironment [31].

## Conclusion

In conclusion, our analysis revealed a considerable overlap between malaria and eBL with regards to their intracellular mechanisms. More comprehensive and integrated cellular, genomic, transcriptomic and proteomic studies will be needed to fully understand the implication of malaria in eBL pathogenesis at molecular level. Verification of the pathways involved in disease pathogenesis and identification of potential targets for therapeutic intervention will provide a solid ground upon which rational drug design and development and objective tailored therapy can be founded.

## Availability of data and materials

The datasets supporting the conclusions of this article are included within the article and its additional files (Supplementary tables)

## Authors’ contributions

Inas Elsayed has participated in all steps of the study. Mutaz Amin and Dr. Muzamil Mahdi Abdel Hamid, heavily participated in the process of re-analysis and re-editing of the paper. Xiaosheng Wang revised the final manuscript. Dr. Mie Rizig has supervised the whole steps of the study and performed the clustering analysis using SOM.

## Acknowledgements

Authors of study acknowledge the help of authors of the three studies from which data has been recruited for this research. Authors especially thank professors; Pier Paolo, Ralf Küppers, and K. Haldar for offering permission to use their data in the study.

## End notes

**a** Gene Ontology (GO) is a collaborative project that aims to reach consistent descriptions of gene products within different databases through development of controlled vocabularies for gene products annotation and development of tools to facilitate their maintenance and use.**^b^** KEGG pathway is a database of manually curated pathway maps expressing knowledge about the molecular interaction and reaction networks of biological systems, such as the cell, the organism and/or the ecosystem.

## Additional files

Table S1. **Malaria and eBL similarly expressed genes - cluster 2 genes.** List of genes in cluster 2 produced by clustering analysis of malaria and eBL gene expression.

Table S2. **Malaria and eBL similarly expressed genes - cluster 5 genes.** List of genes in cluster 5 produced by clustering analysis of malaria and eBL gene expression.

Table S3. **Malaria and eBL similarly expressed genes - cluster 6 genes.** List of genes in cluster 6 produced by clustering analysis of malaria and eBL gene expression.

Table S4. **Malaria and eBL similarly expressed genes - cluster 9 genes.** List of genes in cluster 9 produced by clustering analysis of malaria and eBL gene expression.

Table S5. **list of Genes verified as significantly differentially expressed in eBL.**

Table S6. **list of Genes verified as significantly differentially expressed in *P.falciparum* infected erythroblasts.**

Table S7. **List of pathways enriched in eBL.**

Table S8. **List of pathways enriched in *P.falciparum* infected erythroblasts.**

Table S9.**Functional clusters enriched in eBL.**

Table S10.**Functional clusters enriched in *P.falciparum* infected erythroblasts.**

## List of abbreviations

BL: Burkitt’s Lymphoma
GEO: Gene Expression Omnibus
SOM: Self-Organizing Maps
GO: Gene Ontology
KEGG: Kyoto Encyclopedia of Genes and Genomes
eBL: Endemic Burkitt’s Lymphoma
sBL: Sporadic Burkitt’s Lymphoma
AIDS: Acquired Immunodeficiency Syndrome
EBV: Epstein–Barr virus
NHLs: Non-Hodgkin Lymphomas
GEP: Gene Expression Profile
GC-B lymphocyte: Germinal Centre-B lymphocyte
IGH: Immunoglobulin heavy chain
EBNA-2: Epstein–Barr virus nuclear antigen 2
TNF: Tissue Necrosis Factor
Pf: *Plasmodium falciparum*
HRP-II: histidine rich protein-II
6NANP: Plasmodium falciparum Circumsporozoite Protein
hes-1: hairy and enhancer of split-1
runx3: Runt-related transcription factor 3
JNK: c-Jun N-terminal kinases,
STAT: Signal Transducer and Activator of Transcription
EBNA-1: Epstein–Barr nuclear antigen 1
IFN-γ: Interferon-γ
ART: Artesunate
mRNA: Messenger RNA
Ic50: The half maximal inhibitory concentration
TOGA: total analysis of gene expression
SAGE: serial analysis of gene expression
HG-U133_Plus_2: Affymetrix Human Genome U133 plus 2.0
Array: Array
LIMMA: Linear Models for Microarray Analysis
RMA: Robust Multi-array Analysis Summarization method
DAVID: Database for Annotation, Visualization and Integrated Discovery
MAPK: mitogen-activated protein kinase
CDKs: Cycline dependent Kinases
TGF-β: Transforming Growth Factor-βeta
mTOR: mammalian target of rapamycin

